# Does Communal Breeding Promote an Increase in Social Immunity in Burying Beetles? A Test Case with *Nicrophorus Defodiens*

**DOI:** 10.1101/2021.06.30.450575

**Authors:** M. Suswaram, M. Belk, C. Creighton

## Abstract

Communal breeding is a reproductive system in which more than a single pair of individuals share parental care duties. Burying beetles (genus *Nicrophorus*) breed on small vertebrate carcasses, which is used as a food source for their young. On larger carcasses, burying beetles will breed communally, forming multiple male-female associations. A significant and costly component of parental investment by burying beetles is the preservation of the carcass with secretions containing immune molecules. Because this immune investment is for the benefit of the offspring, the behavior is a form of social immunity. We test the hypothesis that communal breeding in burying beetles evolved as a mechanism to increase the social immune investment on larger carcasses, which are more difficult to preserve. We used *N. defodiens*, a communal breeding burying beetle species to test the hypothesis. There were two experimental treatments wherein, the females either bred communally or non-communally. Our results show that the combined immune activity in the secretions were higher in communally breeding pairs than in the immune contribution of single male-female pairs. However, subordinate females were rarely observed on the carcass, and the level of social immune activity of dominant females was lower than females breeding singly. These data suggest that communal breeding in *N. defodiens* decreases the level of investment in social immunity. Our results demonstrate that the presence of multiple females, which is common under natural conditions, can greatly complicate patterns of social immunity investment in burying beetles.

## Introduction

Communal breeding as a social system is defined primarily by the regular involvement of helpers in the feeding and care of the young (Brown 1978). In some communal breeders, multiple females may breed together sharing duties of parental care of the young (Stacey 1987). Communal breeding behavior is exhibited by a variety of vertebrate and invertebrate species including mammals (Solomon and French 1997), birds (Stacey and Koenig 1990) and insects (Trumbo 1992).

Benefits of communal breeding include shared duties such as obtaining food, defending territories, constructing nests or dens, incubating eggs and defending against predators (Trumbo 1992; Eggert and Sakaluk 2000). In birds and mammals, communal breeding can increase the parent’s reproductive success as the number of helpers increase (Emlen 1991; Dickinson and Hatchwell 2004). Communal breeding can also increase the indirect fitness of helpers, increase the probability of inheriting a territory and improve the chance of survival of the breeder (Woolfenden and Fitzpatrick 1984; Taborsky 1985; Stacey and Ligon 1991). There are several costs associated with communal breeding. For some communally breeding species, females may destroy the eggs of other females (Koenig 1981). Competition with other individuals in a group, may result in a biased reproductive skew among breeding individuals (Scott 1997; Trumbo 1991; Keller and Vargo 1993; Reeves and Ratnieks 1993) and sexually mature individuals may be unsuccessful in producing their own offspring.

Communal breeding can be selected for when the environmental conditions limit the singular breeding attempts of the subordinates – the ‘ecological constraint hypothesis’ (Brown 1987). Communal breeding will also be selected for when it reduces the possibility of takeovers of resources by intra and inter-specific competitors (Eggert and Müller 1992).

Burying beetles are an ideal system to study communal breeding because they exhibit both elaborate biparental care and, in the case of some species, breed communally. Biparental care in burying beetles includes of the preparation and maintenance of small vertebrate carcasses, which is used as a food source by the developing young. Carcass preparation includes removing the fur and feathers from the carcass and covering it with anal and oral exudates that defend the carcass from microbes (Eggert and Müller 1997; Scott 1998). After the young arrive on the carcass, they are fed by the parents with regurgitations of partially broken-down flesh from the carcass (Pukowski 1933).

An important component of parental care in burying beetles is the preservation of carcass with secretions containing immune molecules. This type of immunity where the benefit is directed towards other individuals is referred to as social immunity (Cotter and Kilner 2010). Anal exudates that burying beetles produce for the carcass preservation contain a variety of antimicrobial molecules including lysozyme, antimicrobial peptides and phenoloxidase (Degenkolb et al. 2011). The lysozyme-like activity (LLA) and phenoloxidase (PO) activity appear to trade-off against each other (Cotter and Kilner 2010b). Exudate LLA activity is upregulated and facultatively adjusted upon discovery of a carcass, whereas PO activity is downregulated (Cotter and Kilner 2010b). When a burying beetle is challenged with a personal immunity challenge, social immunity is down regulated, suggesting a tradeoff between the two (Cotter et al. 2010). The LLA levels increase rapidly over two days upon the discovery of a carcass and remain high until the larvae disperse from the carcass. Social immunity is expensive because it diverts resources away from personal immunity functions and shortens life span (Sheldon and Verhulst 1996; Cotter et al. 2010).

On larger carcasses several burying beetle species breed communally with multiple males and females potentially present (Trumbo 1992; Eggert and Sakaluk, 2000). Scott (1994) suggests that in burying beetles, flies and other burying beetles are the major competitors for carcasses and defense against their competitors promotes communal breeding. The evidence supporting this hypothesis is mixed. Eggert and Sakaluk (2000) found no evidence that communal breeding reduces the likelihood of a carcass take over by other burying beetles. Preservation of the carcass and the production of social immunity is expensive. We hypothesize that a female, by tolerating other reproductive females increases the total level of anti-microbial molecules applied to the carcass. This hypothesis predicts that the immune contribution of socially breeding groups should be greater than the immune contribution of single male-female pairs. We test this prediction using the socially breeding burying beetle species *N. defodiens*.

## Materials and Methods

### Burying Beetle Natural History

Burying beetles reproduce on small vertebrate carcasses, which they use as a food resource for themselves and their offspring (Scott 1998). Males and females fight for possession of the carcass, with the largest pair eventually monopolizing it (Wilson and Fudge 1984; Bartlett and Ashworth 1988; Trumbo 1992). *Nicrophorus defodiens* may not bury carrion at all but simply conceals the resource under leaf litter (Trumbo and Bloch 2002; Scott 1998). In *N. defodiens* males remain on average 3.6 days after burial and females remain 7.3 days (Scott and Traniello 1990; Scott 1994) with larvae completing development in 6-7 days after hatching. The offspring emerge from the soil about 25 days later. *N. defodiens* exhibit communal breeding and cooperative brood care on large (generally >80g) carcasses (Eggert and Müller 1992).

### Source of Burying Beetles

Burying beetles used in our experiments were captured in central Wisconsin during May and June 2015, using pitfall traps baited with aged chicken. Wild-caught pairs were placed on a 30-g mouse carcass and allowed to breed to generate the laboratory population. At eclosion, all first generation, laboratory-reared beetles were placed individually in small plastic containers (15.6×11.6×6.7 cm) with *ad libitum* raw chicken liver and maintained on a 14:10 h light/dark cycle. Date of eclosion was designated as day 0 for all subsequent age-based calculations.

### Experimental Design

We randomly assigned females that were 21-25 days old and without previous reproductive experience to one of four treatments. All treatments were represented by 15 replicates. The females were paired with a randomly chosen sexually mature male and placed on a mouse carcass in a large plastic (12 × 9.5 × 5 inches) containers filled with 4 cm of commercially purchased topsoil. There were two control treatments where a single female-male pair was given with a 20g (±1.0 g) or 80g (±1.0 g) carcass and allowed to breed. There were two experimental treatments where two females and one male were given either a 20g (±1.0 g) or 80g (±1.0 g) carcass and allowed to breed. In the experimental treatments, both the females were randomly individually marked with red or green non- toxic, acrylic paint. At the start of each reproductive event, we weighed each female and measured her pronotum width. The design of this experiment is similar to the design followed by Eggert and Müller (1992).

A transparency film (8.25 × 11.7 inches) was marked with three equally spaced elliptical zones. Zone 1 was the closest to the carcass (within 5 inches from the center of the carcass), zone 2 was further away (within 1.5 inches from zone 1) and zone 3 was the farthest from carcass. Observations were made daily from 6 pm to 9 pm. Each day, we recorded the zones in which females were present, number of feeding holes, number of larvae on the carcass, and carcasses were checked daily for the number, instar stage, and dispersal of larvae, as well as death or injury of the female. Dominant female was established as the one closest to the carcass on three or more observations. Each day, from the start of the experiment, the anal exudates were collected from the all the female and a subset of male beetles.

### Phenoloxidase Assay

The fluid samples collected were all tested for the presence of PO activity. For the PO assay 2 μL of sample was added to 100 μL of LPS solution (Sigma-Aldrich L3129) followed by 100 μL of 5 mM L-Dopa (3,4-Dihydroxy-L-phenyl-alanine from Sigma-Aldrich D9628). L-Dopa acts as a substrate for measuring PO (Cotter *et al.* 2010). Samples were then incubated in a BioTek Synergy 2 microplate reader at 30° C and measurements of absorbance taken at 490 nm every minute for an hour. PO activity was expressed as the max change in absorbance of light over the hour.

### Lysozyme-Like Antibacterial Activity

Lysozyme-like activity (LLA) for each fluid sample was measured using a zone of clearance assay. Agar plates containing lyophilized *Micrococcus luteus* were prepared with 1 μL of sample loaded into wells (1.5 mm diameter). For each plate, a control of 1 μL of 1% hen egg white lysozyme (Sigma-Aldrich L6876) was added to one well. The plates were then incubated for 48 h at 27° C. Once removed zone of clearance was measured using Image J software (http://rsweb.nih.gov/ij/index.html).

### Statistical Analysis

To determine the effect of carcass size and social context on immune response and mass change during carcass preparation we used a linear mixed model with repeated measures (MIXED procedure in SAS version 9.4, SAS Institute Inc., Cary, NC, USA). We used three response variables in both males and females – phenoloxidase (PO), lysosome-like activity (LLA), and body mass. Main predictor effects in each model were carcass size (two levels; 20g or 80g), social treatment (three levels for females, nonshared, shared dominant, and shared subordinate; and two levels for males, one or two females), and day of carcass preparation (1-4). Day was used as a repeated measure to quantify changes through time. We included all two-way interactions, and the three-way interaction among main effects. Replicate ID number was used as a random effect to account for the possibility of two females measured simultaneously in some replicates. To determine what part of the test chamber was used by dominant and subordinate females we used a generalized linear model (GENMOD procedure in SAS version 9.4, SAS Institute Inc., Cary, NC, USA). The response variable was zone of the chamber occupied (three levels based on proximity to the carcass). We used a log-link function for the response variable, and main effects were the same as the models used for immune function above. Day was used as a repeated measure to quantify changes through time, and we included all two-way interactions, and the three-way interaction among main effects. A type three analysis was specified to give results similar to the other analyses.

## Results

Dominant and subordinate females differed in the zones in which they were observed during the experiment (Table 1). Dominant females spent 90% of their time closest to the carcass. Subordinate females however, spent most of their time away from the carcass in either zone 2 or zone 3, although they were regularly found on the carcass with multiple feeding holes. Females from both carcass sizes were observed with injuries (χ² = 368.5, df = 18, and P < 0.001), with the subordinate females having far more injuries (Fig. 1). We found no significant difference in body size among males or females across all treatments (Table 2).

**Table 1.** Chi-Square test comparing the different effects of carcass size (carcsize), treatments (trt), and days for zones in which females are present. Statistically significant results (p<0.05) are highlighted in italics.

| Source | DF | Chi-Square | Pr > ChiSq |
| --- | --- | --- | --- |
| Carcsize | 1 | 0.19 | 0.6625 |
| Trt | 2 | 13.23 | **0.0013** |
| Day | 3 | 1.8 | 0.6158 |
| Carcsize*trt | 2 | 4.95 | 0.0841 |
| Carcsize*day | 3 | 2.28 | 0.5163 |
| Trt*day | 6 | 3.48 | 0.7461 |
| Carcsize*trt*day | 6 | 3.33 | 0.7661 |

**Fig. 1.**
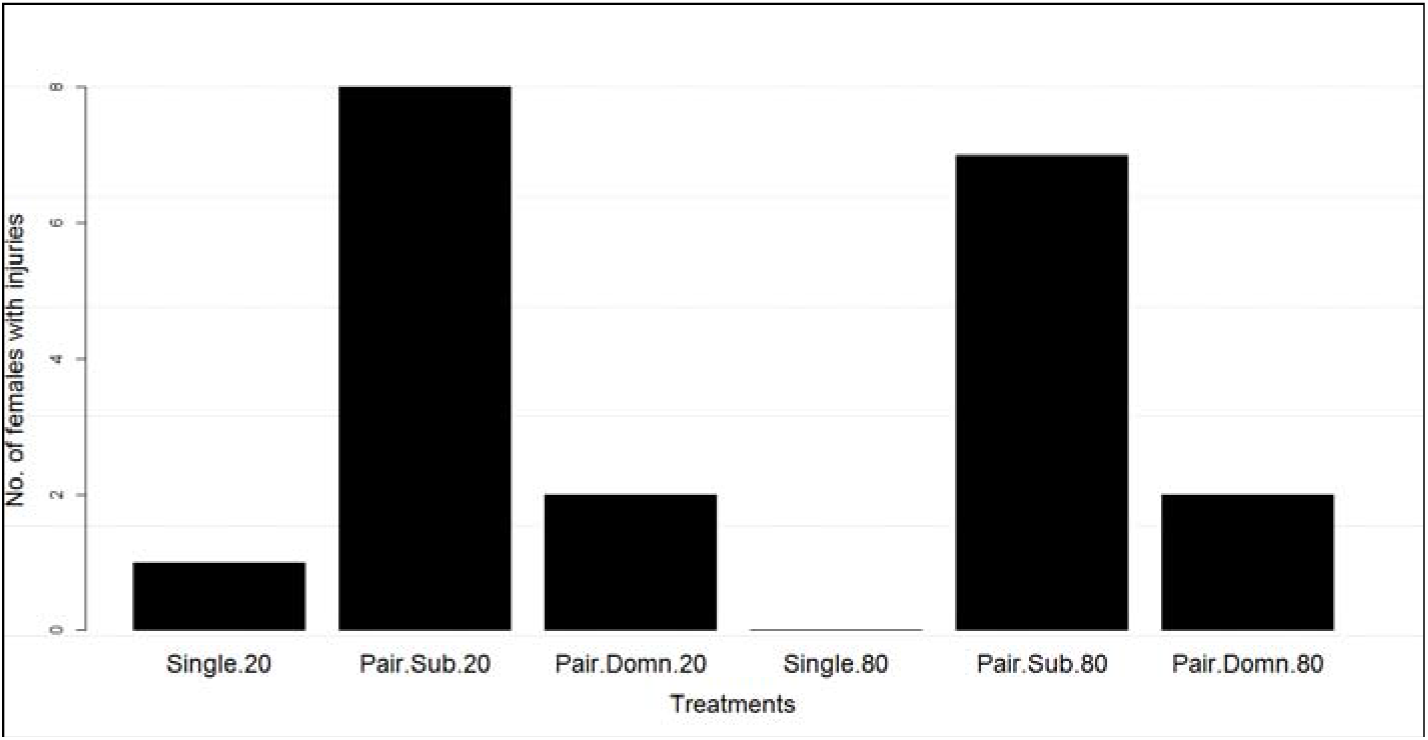
Number of females with injuries in all treatments. Unshared 80g and 20g carcasses represented as Single.80 and Single.20 respectively. Subordinate and dominant females on 20g carcasses and 80g carcasses represented as Pair.Sub.20, Pair.Domn.20, Pair.Sub.80, and Pair.Domn.80 respectively

**Table 2.**
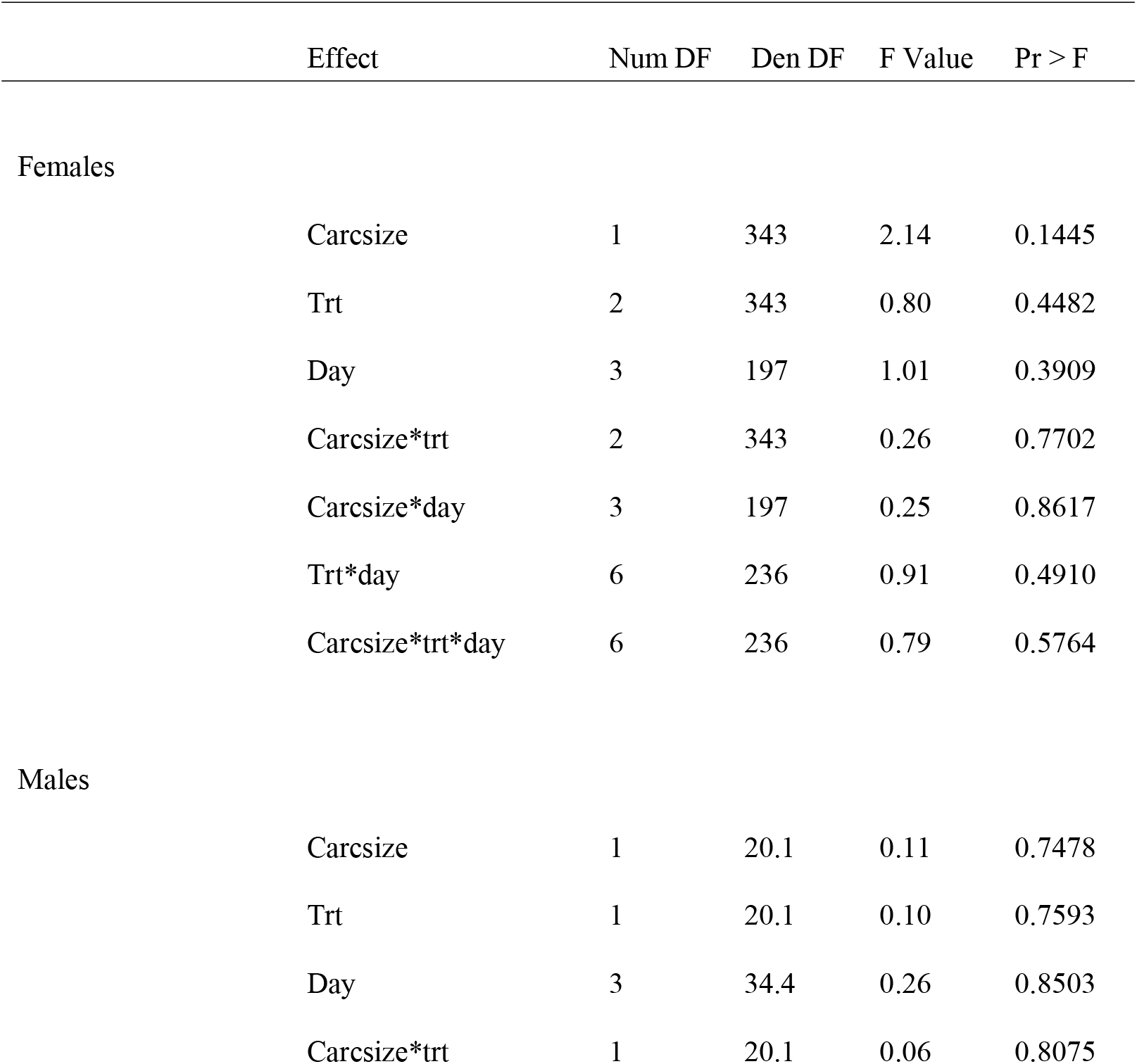
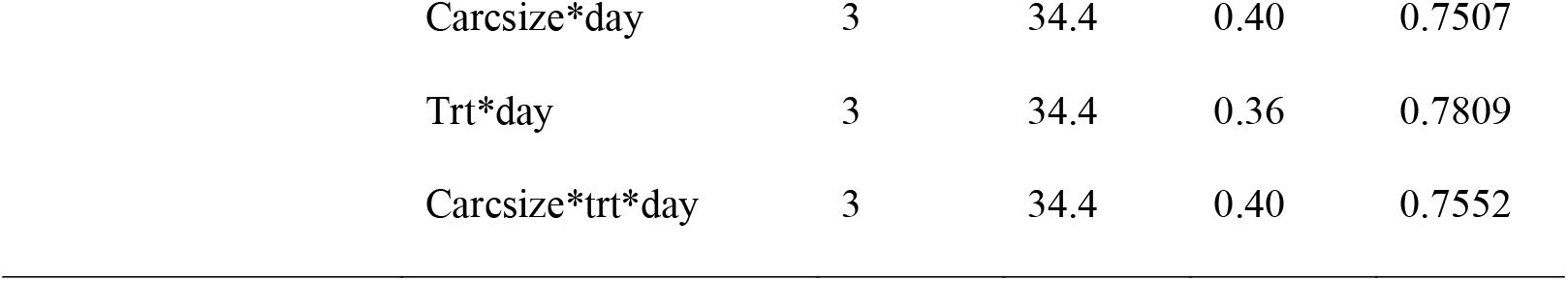
Mixed model ANOVA comparing the different effects of carcass size (carcsize), treatments (trt), and days for body size in females and males.

|  | Effect | Num DF | Den DF | F Value | Pr > F |
| --- | --- | --- | --- | --- | --- |
| Females |  |  |  |  |  |
|  | Carcsize | 1 | 343 | 2.14 | 0.1445 |
|  | Trt | 2 | 343 | 0.80 | 0.4482 |
|  | Day | 3 | 197 | 1.01 | 0.3909 |
|  | Carcsize*trt | 2 | 343 | 0.26 | 0.7702 |
|  | Carcsize*day | 3 | 197 | 0.25 | 0.8617 |
|  | Trt*day | 6 | 236 | 0.91 | 0.4910 |
|  | Carcsize*trt*day | 6 | 236 | 0.79 | 0.5764 |
| Males |  |  |  |  |  |
|  | Carcsize | 1 | 20.1 | 0.11 | 0.7478 |
|  | Trt | 1 | 20.1 | 0.10 | 0.7593 |
|  | Day | 3 | 34.4 | 0.26 | 0.8503 |
|  | Carcsize*trt | 1 | 20.1 | 0.06 | 0.8075 |

|  |  |  |  |  |
| --- | --- | --- | --- | --- |
| Carcsize*day | 3 | 34.4 | 0.40 | 0.7507 |
| Trt*day | 3 | 34.4 | 0.36 | 0.7809 |
| Carcsize*trt*day | 3 | 34.4 | 0.40 | 0.7552 |

Females differed in their level of PO among treatments. There was also a significant carcass size by treatment interaction (Table 3). However, there were no significant effects of carcass size, day or any other interaction (Table 3). PO levels were the highest for the females on unshared 80g and 20g carcasses. Subordinate females on the 80g carcass had the lowest PO levels (Fig. 2a).

**Table 3.**
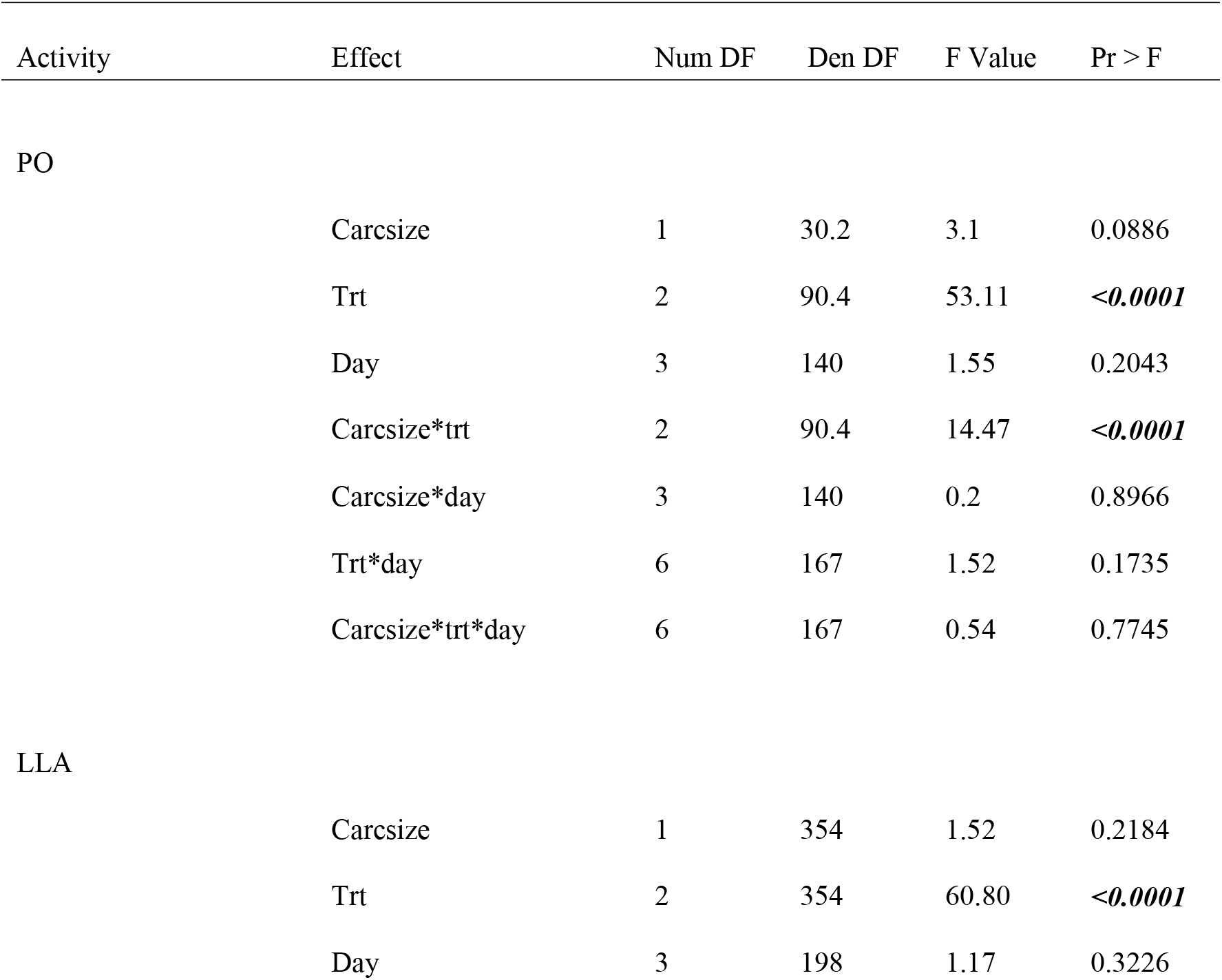
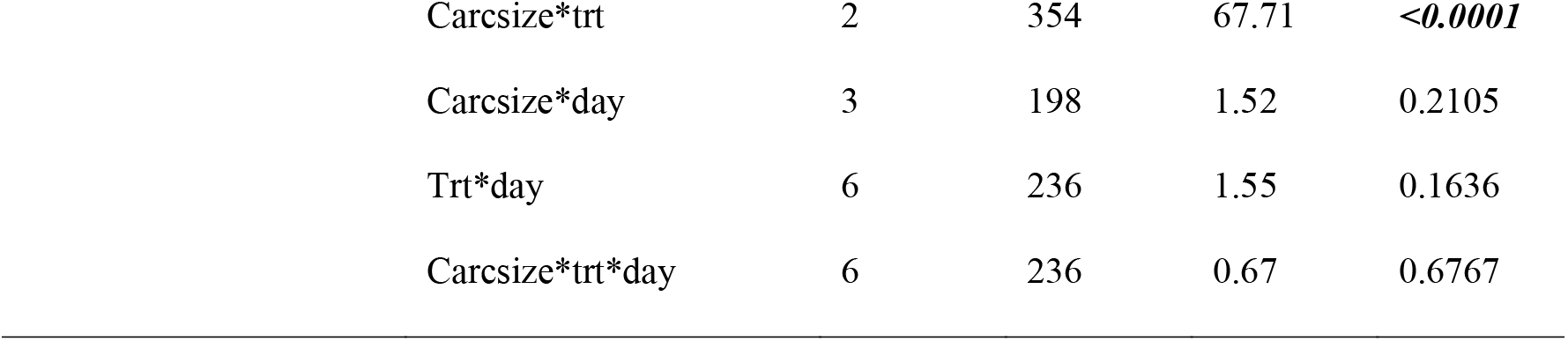
Mixed model ANOVA comparing the different effects of carcass size (carcsize), treatments (trt), and days for PO and LLA in females.

| Activity | Effect | Num DF | Den DF | F Value | Pr > F |
| --- | --- | --- | --- | --- | --- |
| PO |  |  |  |  |  |
|  | Carcsize | 1 | 30.2 | 3.1 | 0.0886 |
|  | Trt | 2 | 90.4 | 53.11 | **<0.0001** |
|  | Day | 3 | 140 | 1.55 | 0.2043 |
|  | Carcsize*trt | 2 | 90.4 | 14.47 | **<0.0001** |
|  | Carcsize*day | 3 | 140 | 0.2 | 0.8966 |
|  | Trt*day | 6 | 167 | 1.52 | 0.1735 |
|  | Carcsize*trt*day | 6 | 167 | 0.54 | 0.7745 |
| LLA |  |  |  |  |  |
|  | Carcsize | 1 | 354 | 1.52 | 0.2184 |
|  | Trt | 2 | 354 | 60.80 | **<0.0001** |
|  | Day | 3 | 198 | 1.17 | 0.3226 |

|  |  |  |  |  |
| --- | --- | --- | --- | --- |
| Carcsize*trt | 2 | 354 | 67.71 | **<0.0001** |
| Carcsize*day | 3 | 198 | 1.52 | 0.2105 |
| Trt*day | 6 | 236 | 1.55 | 0.1636 |
| Carcsize*trt*day | 6 | 236 | 0.67 | 0.6767 |

**Fig. 2.**
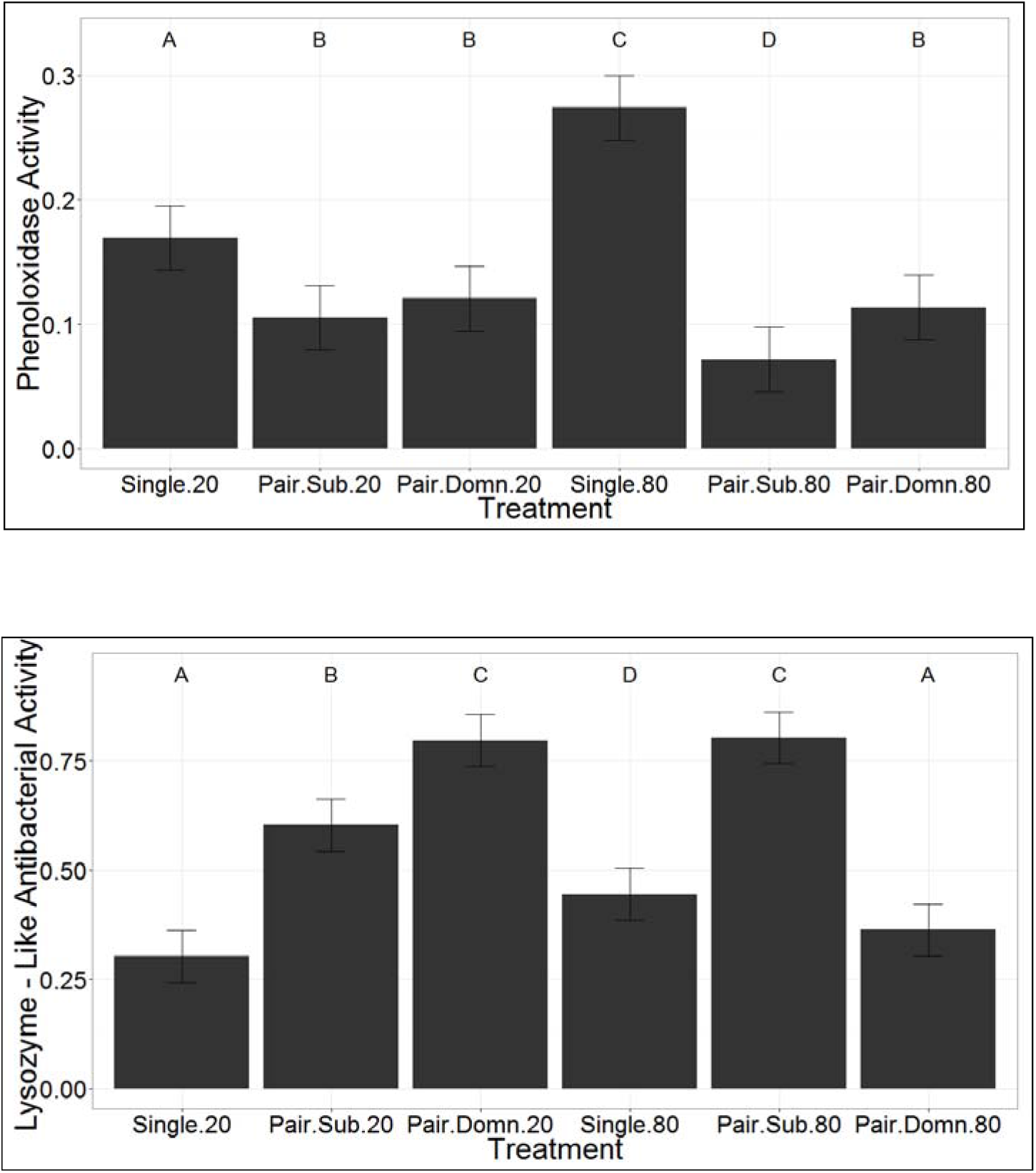
Comparison of the least squared means of (*a*) PO and (*b*) LLA activity for all treatments in females. Letters denote significant differences (*P* < 0.05)

Females differed in their level of LLA among treatments. There was also a significant carcass size by treatment (Table 3). However, there was no significant effects of carcass size, day or any other interaction (Table 3). LLA levels was the highest for subordinate females on the shared 80g carcasses and the dominant females on the shared 20g carcasses. Females on the unshared 20g carcass had the lowest LLA levels (Fig. 2b).

In males, there was a significant effect of carcass size, day, and carcass size by treatment interaction for the levels of PO (Table 4). There was also a significant three-way interaction of carcass size by treatment by day on the levels of PO in males (Table 4). In general, levels of PO decreased across time (Fig. 3a). The males on a shared 20g carcass had significantly higher levels of PO overall (Fig. 3a). The other three treatments significantly differed over time.

**Table 4.** Mixed model ANOVA comparing the different effects of carcass size (carcsize), treatments (trt), and days for PO and LLA in males.

| Activity | Effect | Num DF | Den DF | F Value | Pr > F |
| --- | --- | --- | --- | --- | --- |
| PO |  |  |  |  |  |
|  | Carcsize | 1 | 24.9 | 22.36 | **<0.0001** |
|  | Trt | 1 | 24.9 | 3.35 | 0.0792 |
|  | Day | 3 | 34.9 | 11.56 | **<0.0001** |
|  | Carcsize*trt | 1 | 24.9 | 8.45 | **0.0076** |
|  | Carcsize*day | 3 | 34.9 | 1.34 | 0.3109 |
|  | Trt*day | 3 | 34.9 | 0.85 | 0.4768 |
|  | Carcsize*trt*day | 3 | 34.9 | 3.41 | **0.0280** |
| LLA |  |  |  |  |  |
|  | Carcsize | 1 | 86.7 | 0.45 | 0.5045 |
|  | Trt | 1 | 86.7 | 4.87 | **0.0300** |
|  | Day | 3 | 47.3 | 0.29 | 0.8349 |
|  | Carcsize*trt | 1 | 86.7 | 12.94 | **0.0005** |
|  | Carcsize*day | 3 | 47.3 | 0.02 | 0.9958 |
|  | Trt*day | 3 | 47.3 | 2.54 | 0.0678 |
|  | Carcsize*trt*day | 3 | 47.3 | 1.48 | 0.2328 |

**Fig. 3.**
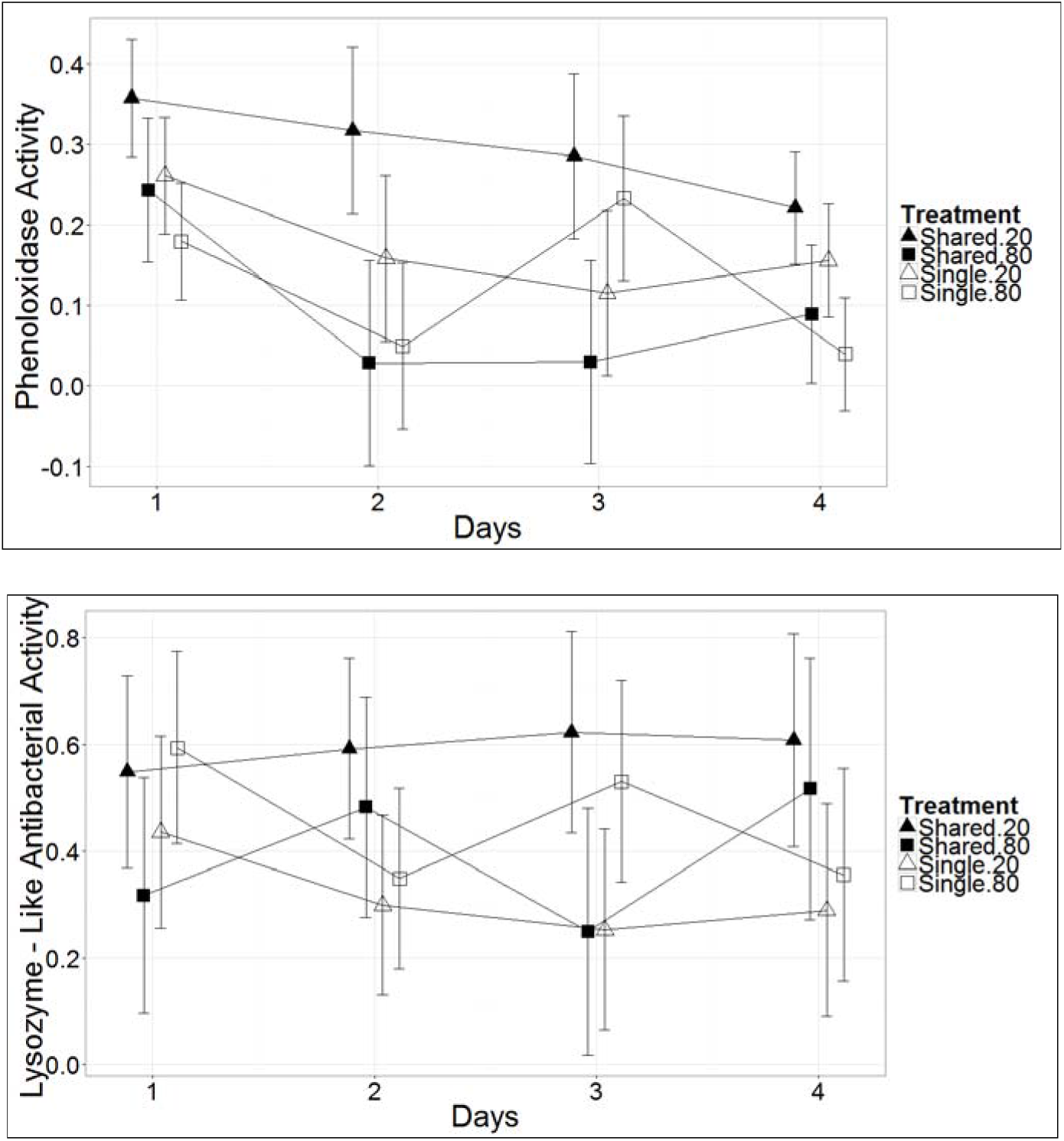
Comparison of the least squared means of (*a*) PO and (*b*) LLA activity for different days and treatments in males

In males, there was a significant effect of treatment, and carcass size by treatment interaction on the levels of PO (Table 4). In general, levels of PO decreased across time (Fig. 3a). The males on a shared 20g carcass had significantly higher levels of LLA overall (Fig. 3a). The other three treatments did not significantly differ over time.

## Discussion

We test the hypothesis that communal breeding in *N. defodiens* increases the combined investment in social immunity on larger carcasses. We tested this by measuring levels of anal secretion in single pair and communally breeding burying beetles. We predicted that the immune contribution of communally breeding pairs will be greater than the immune contribution of single male-female pairs. Previous work has shown that LLA is an important component of the social immune response of burying beetles (Cotter et al. 2010). As predicted, the combined LLA activity of communally breeding females together was much higher than the levels of LLA activity by single females.

However, subordinate females were rarely observed on the carcass although they had elevated LLA levels. With their limited access to the carcass, subordinate females likely made minimal or no contribution to the preservation of the carcass. In addition, LLA levels of dominant, communally breeding females was lower than females breeding singly on the same size carcass. These results suggest that communal breeding in *N. defodiens* decreases the level of investment in social immunity.

A second cost for communally breeding burying beetle females was an increase in the number of injuries. Subordinate females had greater than 50% of the injuries (eight out of observed ten injuries on small carcasses and seven out of nine observed injuries on larger carcasses). Dominant females however, had only two out of ten observed injuries on small carcasses and two out of nine observed injuries on larger carcasses. Even though both the females were breeding, conflict was common. Correspondingly, there was a significant decrease in the levels of PO activity in the anal secretions of females that were communally breeding. These results are consistent with Cotter et al. (2010) who showed that breeding burying beetles given a personal immunity challenge, down regulated their social immunity investment, suggesting a tradeoff between the two.

Dominant and subordinate females differed in the zones in which they were observed in. Dominant females had better access to the carcass and spent most of their time on the carcass. Subordinate females mostly stayed away from the carcass but returned to the carcass multiple times. These results are consistent with the study by Eggert and Müller (1992), who found that on large carcasses fights still occur, but in most cases both females stay on the carcass long enough to provide care for the brood. These results are consistent also with previous research by Eggert and Müller (1992) and Scott and Williams (1993) which showed that in communal associations on large carcasses, a female’s genetic contribution to the joint brood need not be associated with her relative body size and the duration of her stay on the carcass.

The observed pattern of social immunity in males was different than that observed in females. In males, there was a decrease of levels of PO activity in the anal secretions over time, and males breeding communally on 20-g carcasses consistently had higher levels than the other treatments. Males exhibited no clear pattern of LLA levels in the anal secretions over time, except that LLA levels in males breeding jointly on 20g carcasses were consistently higher than the other treatments. There is no strong evidence that communal breeding affects male contribution in any way. However, we interpret our data on male secretion levels with caution due to low sample sizes.

Previous research on social immunity in burying beetles has been done with single female-male pairs. However, multiple male-female associations are commonly observed under natural conditions across a range of carcass sizes (Trumbo 1992; Creighton unpublished data). Our results suggest that social immunity is altered by the changing social structure during a breeding attempt, which can have a significant impact on the ability of burying beetles to preserve their carcass.

